# The *Drosophila* TNF Eiger activates caspase-dependent necrosis when apoptosis is blocked

**DOI:** 10.1101/304964

**Authors:** Mingli Li, Yun Fan

## Abstract

Eiger (Egr), the homolog of the mammalian tumor-necrosis factor (TNF), is the ligand of the c-Jun N-terminal kinase (JNK) stress response signaling pathway in *Drosophila*. Although expression of Egr frequently leads to apoptosis, it has also been implicated in activation of non-apoptotic cell death. However, it is not yet clear how Egr can induce both apoptosis and non-apoptotic cell death, and if so, how such processes are coordinated. Here, we show that expression of Egr in the developing *Drosophila* eye induces apoptosis and non-apoptotic developmental defects, both of which are JNK-dependent. Intriguingly, when apoptotic effector caspases DrICE and Dcp-1 are defective or inhibited, expression of Egr induces necrosis characterized by loss of cell membrane integrity, translucent cytoplasm and aggregation of cellular organelles. Surprisingly, the induction of necrosis depends on the catalytic activity of the initiator caspase Dronc and the input from JNK signaling independently of their roles in apoptosis. Therefore, similar to the mammalian caspase-8, caspases in *Drosophila* also have dual roles in promoting TNF-mediated apoptosis and inhibiting necrosis.

## Introduction

Apoptosis, a major form of programmed cell death, plays crucial roles in elimination of both developmentally unwanted cells and stress-induced damaged cells.^1^ The key factors driving apoptosis are caspases, a family of cysteine proteases. These apoptotic caspases can be further grouped into initiator (or apical) versus effector (or executioner) caspases based on the structure of their N-terminal prodomains and their cleavage substrates.^2^ The initiator caspases have long N-terminal prodomains. Once activated, they further cleave and activate the effector caspases. In contrast, the effector caspases have short prodomains. Once activated, effector caspases cleave a broad spectrum of intracellular substrates to execute cell death. In mammals, caspases can be activated by either intrinsic or extrinsic apoptotic machineries.^3^ Mitochondria are key components in the intrinsic pathway. In response to apoptotic stimuli, cytochrome c is released from mitochondria and binds to the adaptor protein Apaf-1 forming the apoptosome. Caspase-9, an initiator caspase, then interacts with the apoptosome and becomes activated. Activated caspase-9 further cleaves and activates effector caspases such as caspase-3 and –7 leading to apoptosis. Activities of caspase-9, −3 and −7 are also subject to repression by the inhibitor of apoptosis proteins (IAPs). This inhibition can be removed by pro-apoptotic proteins (or IAP antagonists) such as Smac/Diablo and HtrA2/Omi which are also released from mitochondria.^4^

In contrast to the intrinsic pathway, the extrinsic apoptosis pathway is initiated by activation of cell surface death receptors.^3^ Examples of death ligands that can activate death receptors are the tumor necrosis factor (TNF) superfamily, a group of cytokines which was initially discovered because of its anti-tumor activity.^5^ Binding of TNF ligands to their receptors (TNF receptors, TNFRs) promotes TNFR-mediated recruitment of various death-inducing protein complexes depending on the context.^6^ A key component common in these complexes is caspase-8, an initiator caspase. Once activated, caspase-8 can further cleave and activate effector caspases, such as caspase-3 and –7, triggering apoptosis or activate the intrinsic pathway to further enhance the apoptotic response.^7^ Importantly, caspase-8 also cleaves and inactivates two receptor-interacting serine/threonine-protein kinases (RIPKs), RIPK1 and RIPK3, thus preventing activation of necroptosis, another type of programmed cell death (programmed necrosis) which is morphologically distinct from apoptosis.^6, 8^ Therefore, when caspase-8 is deficient or inhibited, activation of TNF and TNFR can lead to necroptosis through RIPK1 and PIPK3. In addition to cell death, functions of TNF family members have also been revealed in immunity, inflammation, cell survival and proliferation.^6^ Many of these functions are mediated by activation of the nuclear factor-κB (NF-κB), a transcription factor controlling multiple target gene expression in a context-dependent manner.^9^ In addition to NF-κB, the TNF signaling can also activate the c-Jun N-terminal kinase (JNK) and the p38 mitogen-activated protein kinase (p38-MAPK) pathways. Both of these pathways are pleiotropic in regulating a variety of cellular processes including cell death, proliferation and differentiation.^10^ It is therefore not surprising that dysregulated TNF-TNFR signaling is associated with numerous pathological situations including cancer and inflammatory diseases.^11^

The core components in the apoptosis pathway are evolutionarily conserved. In *Drosophila*, apoptotic stimuli induce expression of pro-apoptotic proteins Head involution defective (Hid), Reaper (Rpr) and Grim.^12^ These pro-apoptotic proteins act as antagonists to Diap1, the major *Drosophila* IAP, which inhibits activities of both initiator caspases, e.g. Dronc^13, 14^, and effector caspases, e.g. DrICE^15^ and Dcp-1^16^. Once Dronc is released from the inhibition of Diap1, it induces formation of the apoptosome through its interaction with Dark, the *Drosophila* homolog of Apaf-1.^17^ Dronc is then activated. It further cleaves and activates effector caspases such as DrICE and Dcp-1 to induce apoptosis. Compared to mammals, mitochondria are less important in apoptosis in *Drosophila*. However, the pro-apoptotic proteins Hid, Rpr and Grim need to localize to mitochondria to exert their apoptotic functions.^18–20^ Although it is still a subject of debate that the extrinsic apoptosis pathway exists in *Drosophila*, one TNF homolog, Eiger (Egr), has been identified.^21, 22^ Two TNFRs Wengen (Wgn)^23^ and, more recently, Grindelwald (Grnd)^24^ have also been reported. Similar to mammalian TNFs, Egr plays multiple roles in regulating cell death, host defense, pain sensation, nutrient response, tissue growth and regeneration in a context-dependent manner.^25–27^ However, Egr exerts its functions mainly through activation of the JNK pathway in *Drosophila*. For example, expression of Egr under the control of an eye-specific driver *GMR (GMR>egr)* activates JNK and cell death resulting in small adult eyes.^21, 22, 28^ This cell death may not be apoptotic because it cannot be suppressed by P35 (ref.^21^), an inhibitor of the effector caspases DrICE and Dcp-1 (ref.^29^). Paradoxically, *GMR>egr*-induced eye ablation phenotype can be partially rescued by defective apoptosis through reduction of pro-apoptotic gene expression or inhibition of the initiator caspase Dronc.^22, 28^ It is therefore not yet clear whether and how Egr can induce apoptosis and non-apoptotic cell death in parallel.

The core components in the apoptosis pathway are evolutionarily conserved. In *Drosophila*, apoptotic stimuli induce expression of pro-apoptotic proteins Head involution defective (Hid), Reaper (Rpr) and Grim.^12^ These pro-apoptotic proteins act as antagonists to Diap1, the major *Drosophila* IAP, which inhibits activities of both initiator caspases, e.g. Dronc^13, 14^, and effector caspases, e.g. DrICE^15^ and Dcp-1^16^. Once Dronc is released from the inhibition of Diap1, it induces formation of the apoptosome through its interaction with Dark, the *Drosophila* homolog of Apaf-1.^17^ Dronc is then activated. It further cleaves and activates effector caspases such as DrICE and Dcp-1 to induce apoptosis. Compared to mammals, mitochondria are less important in apoptosis in *Drosophila*. However, the pro-apoptotic proteins Hid, Rpr and Grim need to localize to mitochondria to exert their apoptotic functions.^18–20^ Although it is still a subject of debate that the extrinsic apoptosis pathway exists in *Drosophila*, one TNF homolog, Eiger (Egr), has been identified.^21, 22^ Two TNFRs Wengen (Wgn)^23^ and, more recently, Grindelwald (Grnd)^24^ have also been reported. Similar to mammalian TNFs, Egr plays multiple roles in regulating cell death, host defense, pain sensation, nutrient response, tissue growth and regeneration in a context-dependent manner.^25–27^ However, Egr exerts its functions mainly through activation of the JNK pathway in *Drosophila*. For example, expression of Egr under the control of an eye-specific driver *GMR* (*GMR>egr*) activates JNK and cell death resulting in small adult eyes.^21, 22, 28^ This cell death may not be apoptotic because it cannot be suppressed by P35 (ref.^21^), an inhibitor of the effector caspases DrICE and Dcp-1 (ref.^29^). Paradoxically, *GMR>egr*-induced eye ablation phenotype can be partially rescued by defective apoptosis through reduction of pro-apoptotic gene expression or inhibition of the initiator caspase Dronc.^22, 28^ It is therefore not yet clear whether and how Egr can induce apoptosis and non-apoptotic cell death in parallel.

Here, we report that *GMR>egr* primarily induces JNK-dependent apoptosis as well as non-apoptotic developmental defects in the *Drosophila* eye. However, when apoptosis is blocked through inhibition of effector caspases DrICE and Dcp-1, necrosis is induced instead. Intriguingly, loss of one copy of the gene encoding the initiator caspase Dronc suppresses necrosis but not Egr-induced apoptosis. Moreover, activation of regulated necrosis requires the catalytic activity of Dronc as well as the additional input from the JNK signaling pathway. Therefore, caspases are crucial in regulating TNF-triggered necrosis in *Drosophila* which is analogous to its mammalian counterparts.

## Results

### The cleaved Dcp-1 antibody is a specific marker for activated effector caspases DrICE and Dcp-1 in *Drosophila*

To determine whether *GMR>egr* induces apoptosis and/or non-apoptotic cell death, we first sought to identify a marker that specifically recognizes activated effector caspases, e.g. cleaved DrICE and Dcp-1, in *Drosophila* because antibodies recognizing the cleaved human caspase-3 are not specific to cleaved DrICE and Dcp-1.^30^ The recently developed cleaved Dcp-1 (Asp216) antibody (referred to as cDcp1) from Cell Signaling Technology is a polyclonal antibody raised against the large 22 kDa fragment of cleaved Dcp-1. Although this antibody has been increasingly used to label apoptosis in *Drosophila* ^31^, a detailed characterization of its specificity has not been carried out. We therefore addressed this by using *GMR-hid*, a transgene leading to induction of apoptosis specifically in the developing *Drosophila* eye.^32^ In late 3^rd^ instar eye discs, compared to wild type where apoptosis occurs at a very low level (Fig.2e), *GMR-hid* induces two waves of apoptotic cells as indicated by TUNEL, an assay detecting DNA fragmentation a hallmark of apoptotic cells (Fig.1a).^33^ cDcp1 antibodies also recognize these two apoptotic waves in *GMR-hid* as expected (Fig1b). To determine whether cDcp1 only detects the cleaved Dcp-1, we analyzed its staining in *dcp-1* null mutants which does not obviously affect GMR-hid-induced apoptosis (Fig. 1c). Intriguingly, cDcp1-labeling of two apoptotic waves persists (Fig.1d) suggesting at least one other apoptotic protein is recognized by cDcp1. In addition to Dcp-1, DrICE is the second major effector caspase mediating apoptosis in somatic tissues including eye discs.^34, 35^ Moreover, DrICE and Dcp-1 share the same cleavage site for their activation.^30, 36^ It is therefore possible that cDcp-1 also recognizes the cleaved DrICE. To examine this, we labeled *GMR-hid* discs with cDcp1 in *drICE* null mutants. Although apoptosis is almost completely lost in *drICE* mutants as indicated by TUNEL (Fig.1e), cDcp1 detects a relatively low level of signals in the *GMR* domain (Fig.1f) which are presumably the cleaved Dcp-1 proteins. Importantly, cDcp1-and TUNEL-labeling is lost in *dcp-1; drICE* double null mutants (Fig.1g,h). Therefore, cDcp1 recognizes both cleaved Dcp-1 and cleaved DrICE.

**Figure 1.**
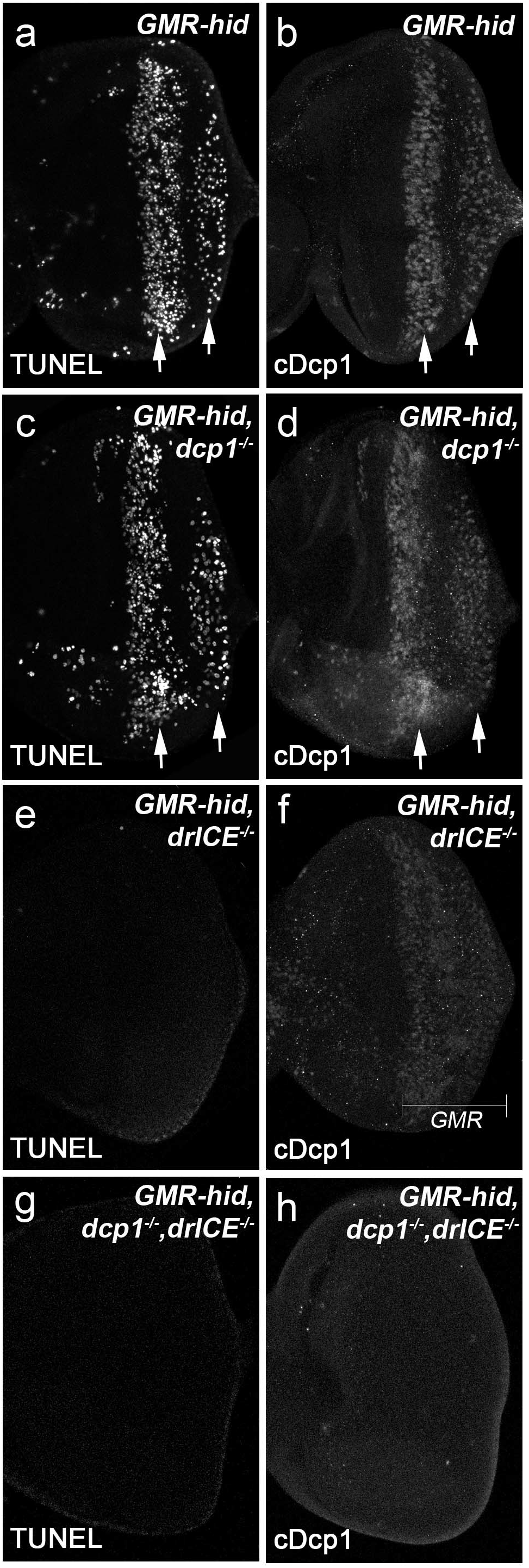
The cDcp1 antibody recognizes the cleaved DrICE and Dcp-1 in *Drosophila*. Late 3^rd^ instar larval eye discs labeled with either the TUNEL assay (a,c,e,g) or the cDcp1 antibodies (b,d,f,h). (a, b) Expression of *hid* under the control of *GMR*(*GMR-hid*) induces two apoptotic waives indicated by either TUNEL (a, arrows) or cDcp1 (b, arrows) staining. (c, d) *GMR-hid*-induced two apoptotic waves are recognized by both TUNEL (c) and cDcp1 (d) in *dcp-1* null mutants. (e, f) *GMR-hid*-induced apoptosis is blocked in *drICE* null mutants as indicated by no TUNEL labeling (e). However, cDcp1 recognizes a low level of signals in the whole *GMR* domain in the same genetic background (f). (g, h) No *GMR-hid*-induced signals were detected by either TUNEL (g) or cDcp1 (h) in *drICE* and *dcp-1* double null mutants.

**Figure 2.**
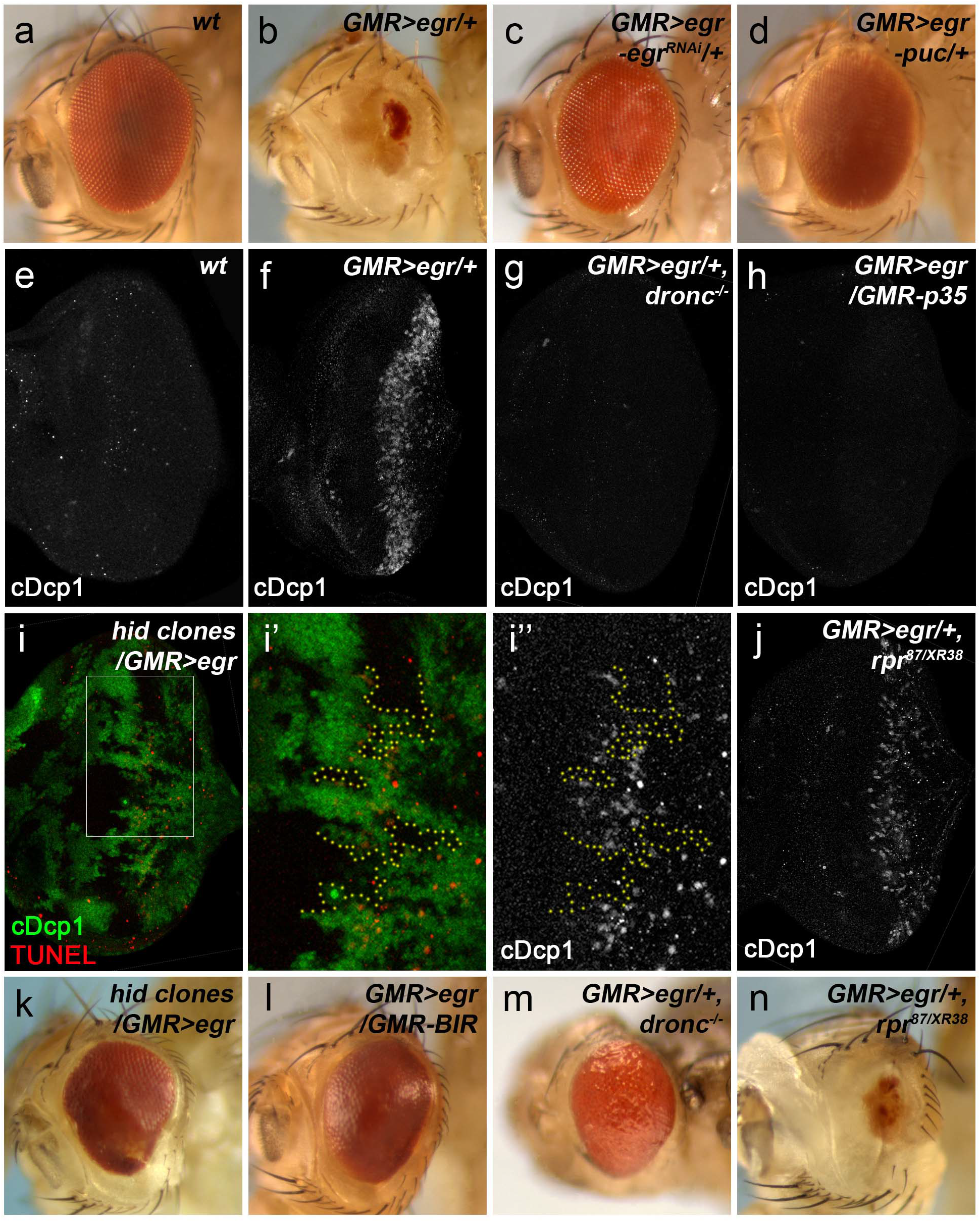
*GMR>egr* induces apoptosis through Hid in the *Drosophila eye*. (a-d) Adult eye images. Compared to wild type (a), *GMR>egr* induces a strong eye ablation phenotype (b). This phenotype is completely suppressed by an RNAi knockdown of *egr*(*egr^RNAi^*, c) or expression of *pucker* (*puc*, d), a negative regulator of JNK. (e-j) Late 3^rd^ instar eye discs labeled with cDcp1 (a-h,j) or cDcp1 and GFP (i-i’’). Compared to wild type (e), expression of egr under the control of GMR (GMR-GAL4/UAS-egr, *GMR>egr*) induces massive apoptosis indicated by cDcpl labeling (f). This apoptosis is suppressed in *dronc* null mutants (g) or by expression of p35 (GMR-p35, h). In *GMR>egr* discs with *hid* mutant clones marked by lack of GFP (i), apoptosis is blocked in the clones (highlighted by yellow dotted lines in i’ and i’’ which are enlarged images of the outlined 87/XR38a area in i). In contrast, *rpr* mutants (*rpr^87/XR38a^*, combination of a deletion and a null mutant of *rpr*) do not suppress *GMR>egr*-induced apoptosis (j). (k-n) Adult eye images. *hid* mutant clones (k), expression of a RING domain-deleted, therefore stabilized, form of Diapl (GMR-BIR, l) or *dronc* null mutants (pharate adults were dissected out of the pupal cases, m) strongly suppress *GMR>egr*-induced eye ablation 87/XR38a phenotype. In contrast, *rpr* mutants (*rpr^87/XR38a^*) do not suppress small eyes induced by *GMR>egr* (n).

### Expression of Eiger induces strong apoptosis through the canonical apoptosis pathway and the pro-apoptotic gene *hid*

*GMR>egr* induces a strong eye ablation phenotype (compare 2b to 2a).^21, 22^ As controls, we validated that *GMR>egr*-induced small eyes are completely suppressed by knocking down *egr* itself through RNAi (Fig.2c) or expressing *pucker (puc)* (Fig.2d), a negative regulator of JNK.^37, 38^. Therefore, *GMR>egr*-induced eye ablation is indeed due to expression of *egr* and activation of JNK. To assess whether *GMR>egr* induces apoptosis, we used cDcp1 to label 3^rd^ instar larval eye discs in which the *GMR* promoter is expressed. Compared to wild type (Fig.2e), a strong wave of cDcp1-labeling was observed in *GMR>egr* discs (Fig.2f). To further confirm that apoptosis is induced by *GMR>egr* genetic analyses on key components of the canonical apoptosis pathway were conducted. Loss of Dronc, the major initiator caspase mediating apoptosis in *Drosophila* ^39, 40^, or expression of P35, an inhibitor of activated DrICE and Dcp-1 (ref.^29^), completely blocks cDcp1 signals induced by *GMR>egr* (Fig.2g,h). Consistently, although *dronc* null mutants are mostly pharate adult lethal, they show suppressed *GMR>egr* eyes (compare 2m to 2b) when they are dissected out of the pupal cases. Furthermore, expression of a RING domain-deleted (therefore stabilized) form of Diap1 (BIR), the apoptosis inhibitor upstream of Dronc ^41^, also strongly suppresses *GMR>egr*-induced small eyes (compare Fig.2l to 2b). Therefore, *GMR>egr* induces massive apoptosis in the developing *Drosophila* eye.

To identify which pro-apoptotic genes mediate *GMR>egr*-induced apoptosis, we first examined expression of *hid* and *rpr* by using their reporters.^42^ Compared to the control, expression of *hid*, but not *rpr*, shows significant increase (Supplementary Fig.S1) in response to *egr* expression. This is consistent with a previous report.^28^ To determine whether *hid*, but not *rpr*, is required for *GMR>egr*-induced apoptosis, mutant analyses were performed. While loss of *rpr* by using a combination of a deletion ^43^ and a null mutant of *rpr* (ref.^44^), *rpr87/XR38*, does not significantly affect the level of *GMR>egr*-induced apoptosis (Fig.2j), *GMR>egr*-induced apoptosis is lost in *hid* mutant clones (Fig.2i-i’’). Consistently, *GMR>egr*-induced small eyes are suppressed by *hid* mosaics (Fig.2k), but not by *rpr* mutants (Fig.2n). Taken together, expression of Egr activates apoptosis through the pro-apoptotic gene *hid* in the developing *Drosophila* eye.

### Expression of Eiger also induces JNK-dependent non-apoptotic defects

Intriguingly, unlike expression of *puc* (Fig.2d), *loss-of-dronc* or expression of Diap1 *(GMR>BIR)* does not completely restore *GMR>egr* eyes back to normal indicated by their glassy appearance suggesting irregular ommatidial patterning (Fig.2i,m). Thus expression of Egr in the eye can induce defects other than apoptosis. These defects appear to depend on JNK activity. Similar to expression of *puc*, expression of a dominant negative form of *bsk (bsk^DN^, bsk=Drosophila JNK)* or hemizygous mutants of *Takl (Tak1^2527^*, null mutant ^45^), an upstream kinase of JNK, almost completely rescue *GMR>egr*-induced adult eye defects (Fig.3a,b). Therefore, in addition to apoptosis, *GMR>egr* can also induce JNK-dependent non-apoptotic defects (Fig.3c).

**Figure 3.**
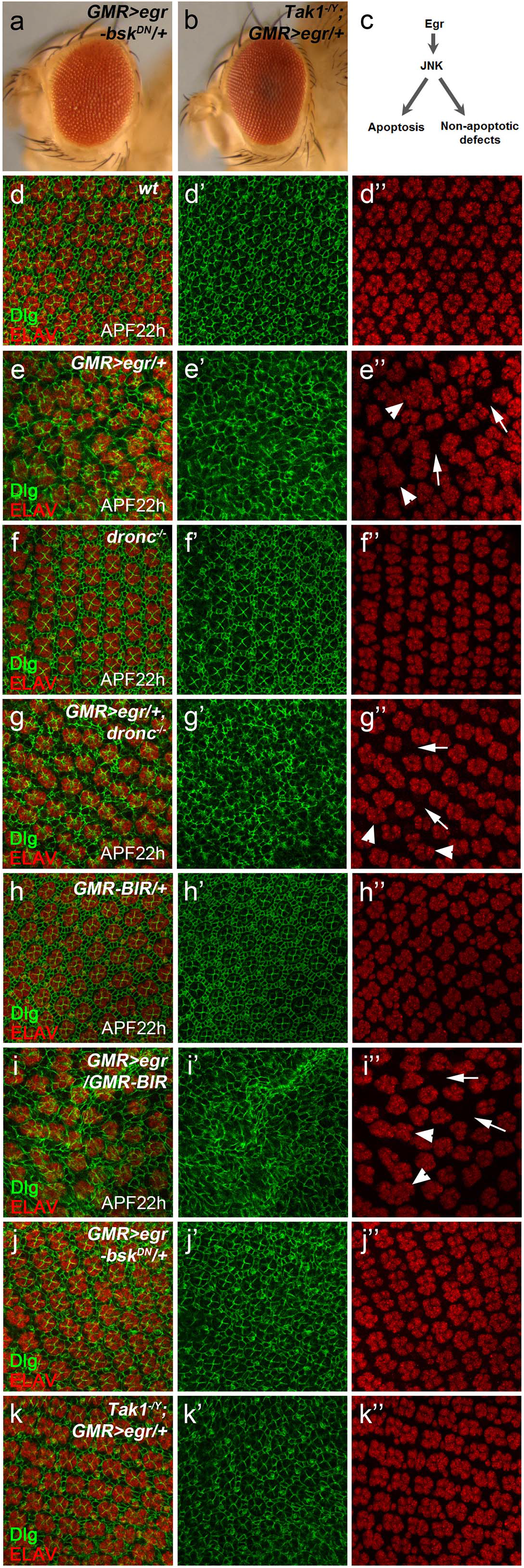
*GMR>egr* induces non-apoptotic, but JNK-dependent, defects in the *Drosophila eye*. (a, b) Adult eye images. Expression of a dominant negative form of *bsk* (*bsk^DN^, bsk=Drosophila JNK*) (a) or hemizygous mutants of *Takl* (b), an upstream kinase of JNK, almost completely suppresses *GMR>egr*-induced eye ablation phenotype. (c) A diagram showing the JNK signaling induced by Egr can lead to both apoptosis and non-apoptotic defects in the developing eye. (d-k’’) APF22h pupal eye discs labeled with a cellular membrane maker Dlg (green in d-k and d’-k’) and a neuronal marker ELAV (red in d-k and d’’-k’’). In wild type discs (d-d’’), ommatidia, each is composed of eight photoreceptor neurons as indicated by ELAV, and interommatidial cells, as indicated by Dlg, are well patterned. In contrast, defects in cellular organization were observed in *GMR>egr* discs (e-e’’). Examples of these defects such as ommatidial fusion (arrowheads in e’’,g’’,i’’) and increased interommatidial spacing (arrows in e’’,g’’,i’’) are highlighted. *dronc* null mutants (f-f”, *dronc^−/−^*) or expression of a stabilized form of Diapl (h-h’’, *GMR-BIR*) neither alter the ommatidial patterning in wild type eye discs by themselves (compare f-f” and h-h’’ to d-d’’) nor suppress the irregular ommatidial organization in *GMR>egr* discs (compare g-g’’ and i-i’’ to e-e’’). In contrast, expression of a dominant-negative form of JNK (*bsk^DN^*, j-j’’) or hemizygous mutants of *Takl* (k-k’’) strongly suppresses the defects in cellular organization induced by *GMR>egr*.

To further characterize *GMR>egr*-induced non-apoptotic defects at the cellular level, we examined ommatidial organization in pupal eye discs which is a sensitive readout of defects in eye development. Developing pupal eye discs are composed of well-patterned ommatidia (Fig.3d,d’’, ELAV labelling) and interommatidial cells (Fig.3d,d’). At 25°C, developmental apoptosis occurs at around 28h after puparium formation (APF28h) to remove extra interommatidial cells in eye discs (Supplemental Fig.S2a-a’’).^46, 47^ However, no apoptosis is observed at APF22h (Fig.4a,a’). We therefore performed our analysis of *GMR>egr* at APF22h to avoid developmental apoptosis. Compared to wild type eye discs where ommatidia and interommatidial cells are well-organized (Fig.3d-d’’), *GMR>egr* leads to irregular organization of ommatidia and interommatidial cells (Fig.3e-e’’) with both increased interommatidial spacing (Fig.3e’’, arrows) and ommatidial fusion (Fig.3e’’, arrowheads) observed. These *GMR>egr*-induced defects persist when apoptosis is inhibited in *dronc* null mutants (Fig.3g-g’’) or by expression of Diap1 (Fig.3i-i’’). As controls, at APF22h, ommatidial organization is not affected by loss-of-dronc (Fig.3f-f”) or expression of Diap1 (Fig.3h-h’’) alone. In contrast, *GMR>egr*-induced ommatidial organization defects are strongly suppressed by expression of *bsk^DN^* (Fig.3j-j’’) or in hemizygous mutants of *Tak1*(Fig.3k-k’’). Altogether, these data indicate that *GMR>egr* induces both apoptosis and non-apoptotic, but JNK-dependent, defects in the developing *Drosophila* eye (Fig.3c).

**Figure 4.**
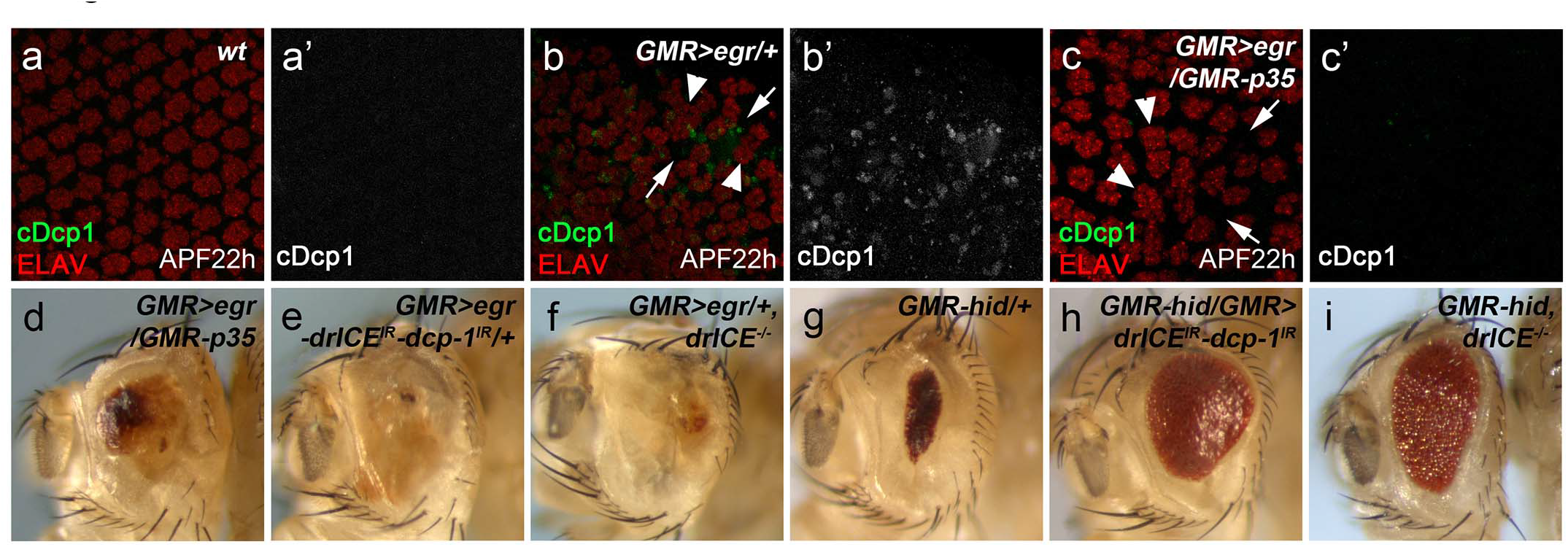
Non-apoptotic cell death is induced in *GMR>egr* when effector caspases DrICE and Dcp-1 are inhibited. (a-c’) APF22h pupal discs labeled with cDcpl (green in a,b,c and grey in a’,b’,c’) and ELAV (red in a,b,c), a neuronal marker. At this stage, no apoptotic cells were detected in wild type discs (cDcpl, a and a’). Ommatidia are also well patterned (ELAV, a). In contrast, strong apoptosis were detected at APF22h in *GMR>egr* eye discs (cDcpl, b and b’). Defects in cellular organization indicated by increased interommatidial spacing (arrows, b) and ommatidial fusion (arrowheads, b) were observed. Although *GMR>egr*-induced apoptosis is almost completely blocked by expression of P35 (cDcpl, c and c’), the irregular ommatidial organization (c, arrows and arrowheads) is not rescued. (d-i) Adult eye images. Expression of P35 (*GMR-p35*, d), RNAi knockdown of *drICE* and *dcp-1* (e), or *drICE* null mutants (f) do not or only slightly suppress *GMR>egr*-induced eye ablation phenotype (compare 4d,e,f to 2b). In contrast, *GMR-hid*-induced small eyes (g) can be suppressed by RNAi knockdown of *drICE* and *dcp-1* (h) or *drICE* null mutants (i).

### Inhibition of effector caspases in *GMR>egr* induces non-apoptotic cell death

Another intriguing observation that has been reported previously is that, unlike *dronc* mutants or expression of Diap1, expression of P35, an inhibitor of effector caspases, cannot or only slightly suppress *GMR>egr*-induced eye ablation phenotype (compare Fig.4d to Fig.2b, quantified in Fig.5i). ^21,22,48^ One possibility is that expression of P35 may not be sufficient to block *GMR>egr*-induced apoptosis because P35 is a pseudo-substrate of effector caspases.^29, 49^ We looked into this possibility by using cDcp1 to label apoptotic cells. As a control, cDcp1 detects developmental apoptosis in pupal discs at APF28h (Supplementary Fig.S2). We therefore examined eye discs at APF22h to avoid developmental apoptosis. Compared to the wild type discs where no apoptosis is detected at APF22h (Fig.4a,a’), *GMR>egr* induces massive apoptosis and defective ommatidial organization (Fig.4b,b’). Notably, *GMR>egr*-induced apoptosis, but not ommatidial mis-organization, is almost completely blocked by expression of P35 (Fig.4c,c’). However, unlike *dronc* mutants or expression of Diap1, expression of P35 in *GMR>egr* still results in small eyes although it suppresses *GMR>egr-induced* apoptosis (compare Fig.4d to Fig.2l,m). Importantly, this is not due to any unspecific effects induced by P35 itself because double RNAi knockdown of *drICE* and *dcp-1* or using *drICE* mutants does not suppress, or even enhance, *GMR>egr*- induced eye ablation phenotype (Fig.4e,f). Both knockdown of *drICE* and *dcp-1* and loss-of-*drICE* can suppress *GMR-hid-induced* apoptosis confirming these RNAi lines and mutants are functional (Fig.4g-i). Taken together, these data suggest that inhibition of effector caspases, in particular DrICE, may lead to non-apoptotic cell death, while it suppresses apoptosis in *GMR>egr*.

**Figure 5.**
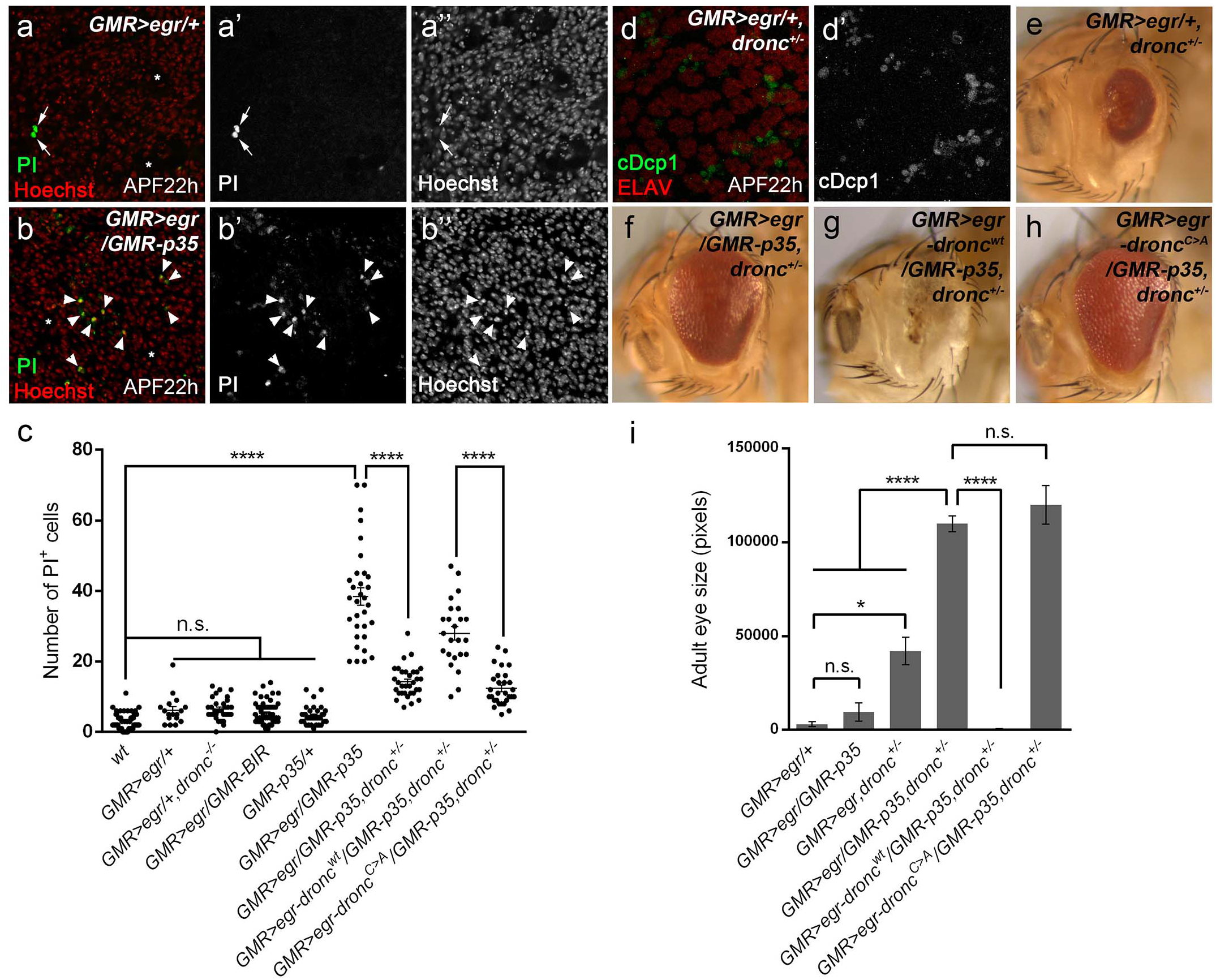
*GMR>egr* induces Dronc-dependent non-apoptotic cell death when effector caspases DrICE and Dcp-1 are inhibited. (a-b’’) APF22h pupal eye discs labeled with Propidium Iodide (PI, green in a,b and grey in a’,b’) and Hoechst (red in a,b and grey in a’’,b’’). In *GMR>egr* discs, PI detects a background level of signals (arrows, a-a’’) which often do not co-localize with the Hoechst labeling, a nucleus marker. In contrast, expression of P35 (*GMR>egr/GMR-p35*) results in a strong increase of PI-positive nuclei, majority of which are also Hoechst-positive (arrowheads, b-b’’). Asterisks indicate irregular cellular spacing in both *GMR>egr* and *GMR>egr/GMR-p35* discs. (c) Quantification of PI-positive cell numbers in APF22h pupal eye discs of various genetic backgrounds as indicated. One-way ANOVA with Bonferroni multiple comparison test was used to compute p-values. Asterisks indicate statistically significant changes (p<0.0001). A background low level of PI-labeling was observed in both wild type and *GMR>egr discs*. This low PI-labeling in *GMR>egr* is not increased in *dronc* mutants or by expression of a stabilized form of Diap-1 (*GMR-BIR*). PI-labeling is also low in *GMR-p35* discs. In contrast, strong PI-labeling was observed in *GMR>egr/GMR-p35* discs. This PI-labeling is largely suppressed in *dronc* heterozygous mutants (*GMR>egr/GMR-p35, dronc^+/−^*). In this background, further expression of a wild type form of Dronc (*GMR>egr-dronc^wt^/GMR-p35, dronc^+/−^*), but not a catalytic site-mutated form of Dronc (*GMR>egr-dronc^C>A^/GMR-p35, dronc^+/−^*), is sufficient to restore PI-labeling. (d, d’) APF22h pupal eye discs labeled with cDcp1 (green in d and grey in d’) and ELAV (red in d). Loss of one copy of *dronc* does not or only slightly suppress *GMR>egr*-induced apoptosis. (e-h) Adult eye images. Although loss of one copy of *dronc* only slightly suppresses *GMR>egr*-induced small eye phenotype (compare 5e to 2b), it strongly rescues the eye ablation phenotype induced by *GMR>egr/GMR-p35* (compare 5f to 4a). This rescue is neutralized by expression of a wild type form of Dronc (g), but not a catalytic site-mutated form of Dronc (h). (i) Quantification of the average adult eye size (mean±SD) of various genetic backgrounds as indicated. One-way ANOVA with Bonferroni multiple comparison test was used to compute p-values. Asterisks indicate statistically significant changes (P<0.1 or P<0.0001 as indicated by one or four asterisks). Suppression of *GMR>egr* by expresssion of P35 is not statistically significant. Heterozygous *dronc* mutants only weakly suppress *GMR>egr*-induced small eyes (*GMR>egr*/+, *dronc^+/−^*). But they strongly suppress *GMR>egr/GMR-p35*-induced eye ablation phenotype (*GMR>egr/GMR-p35, dronc^+/−^*). In this background, further expression of a wild type form of Dronc (*GMR>egr-droncwt/GMR-p35, dronc^+/−^*), but not a catalytic site-mutated form of Dronc (*GMR>egr-dronc^c>A^/GMR-p35, dronc^+/−^*), is sufficient to restore the eye ablation phenotype.

### The initiator caspase Dronc mediates Eiger-induced necrosis when apoptosis is blocked

To further characterize the non-apoptotic cell death induced by *GMR>egr* when effector caspases are inhibited, we performed labeling with propidium iodide (PI), a dye that enters the cells and binds to their nucleic acids when the cell membrane integrity is disrupted. Therefore, PI has been used to label necrotic cells, characterized by membrane rupture, in both *Drosophila* eye and wing tissues (Supplementary Fig.S3a,b)^50, 51^. Interestingly, no PI-positive cells were detected in late 3^rd^ instar larval *GMR>egr* discs with or without expression of P35 (Supplementary Fig.S3c,d) suggesting the non-apoptotic cell death may not occur immediately following apoptosis inhibition. We then examined eye discs in the pupal stage of APF22h to allow more time for non-apoptotic cell death to occur. In *GMR>egr* a very low level of PI-labeling was observed (Fig.5a,a’). Notably, most of these PI signals do not co-localize with a DNA marker Hoechst (Fig.5a,a’’, arrowheads) suggesting that such a low level of PI-labeling may be unspecific. This is further supported by a comparable low level of PI signals observed in wild type discs (quantified in Fig.5c) under the same experimental condition. Therefore, only a low level of, probably unspecific, PI-labeling can be detected in wild type or *GMR>egr* pupal discs. In contrast, expression of P35 strongly induces PI-labeling in *GMR>egr* pupal discs (Fig.5b,b’). Importantly, the majority of these PI signals co-localizes with Hoechst-positive nuclei (Fig.5b,b’’, arrowheads) suggesting that they specifically label membrane-compromised cells. As a control, expression of P35 alone does not induce PI-labeling (Fig.5c). Altogether, these data suggest that a factor(s) upstream of effector caspases may trigger necrosis when apoptosis is blocked by P35. We therefore examined PI-labeling in *GMR>egr* discs in a background of *dronc* mutants or with expression of the stabilized Diap1 *(GMR-BIR)* as they both suppresses *GMR>egr*-induced apoptosis upstream of effector caspases (see Fig.2l,m). Only background levels of PI-labeling were detected in these conditions (Fig.5c). Strikingly, loss of one copy of *dronc*, although it only weakly affects *GMR>egr*-induced apoptosis (Fig.5d,d’) and eye ablation phenotype (Fig.5e,i), it strongly suppresses PI-labeling induced in *GMR>egr/GMR-p35* (Fig.5c). Consistently, *dronc* heterozygous mutants together with expression of P35 strongly suppress *GMR>egr*-induced eye ablation phenotype (Fig.5f,i). Therefore, Dronc is a key component mediating PI-positive cell death when *GMR>egr*-induced apoptosis is blocked by expression of P35.

It is known that Dronc, the initiator caspase, cleaves its downstream effector caspases DrICE and Dcp-1 to activate apoptosis in Drosophila.^13, 36^ We thus examined whether the catalytic activity of Dronc is required for non-apoptotic cell death induced in *GMR>egr/GMR-p35* eye discs. To do this, either a wild type Dronc *(droncwt)* or a catalytic site-mutated form of Dronc *(droncC>A)* transgene was used.^13^ As expected, expression of the wild type Dronc restores PI-positive cell death (Fig.5c), and therefore eye ablation (Fig.5g,i), in *GMR>egr/GMR-p35*; dronc^+/−^ animals. However, expression of *droncC>A* in the same background does not have these effects (Fig.5c,h,i). This suggests that the catalytic activity of Dronc is required for Egr-induced non-apoptotic cell death when apoptosis is blocked.

To further determine the type of non-apoptotic cell death induced in *GMR>egr/GMR-p35*, we performed the Transmission Electron Microscopy (TEM) analysis on APF22h pupal eye discs. Compared to the wild type control (Fig.6a), cells with typical apoptotic features ^8^ such as high-electron-density chromatin condensation and apoptotic bodies were frequently observed in *GMR>egr* discs (Fig.6b,b’). In contrast, cells with typical necrotic features were observed in *GMR>egr/GMR-p35* eye discs (Fig.6c-c’’). These cells often have translucent cytoplasm, mal-shaped or unidentifiable nuclei, and aggregation of endoplasmic reticulum and other cellular organelles.^8, 52^ Therefore, it appears that necrosis is induced in *GMR>egr/GMR-p35* pupal eye discs.

**Figure 6.**
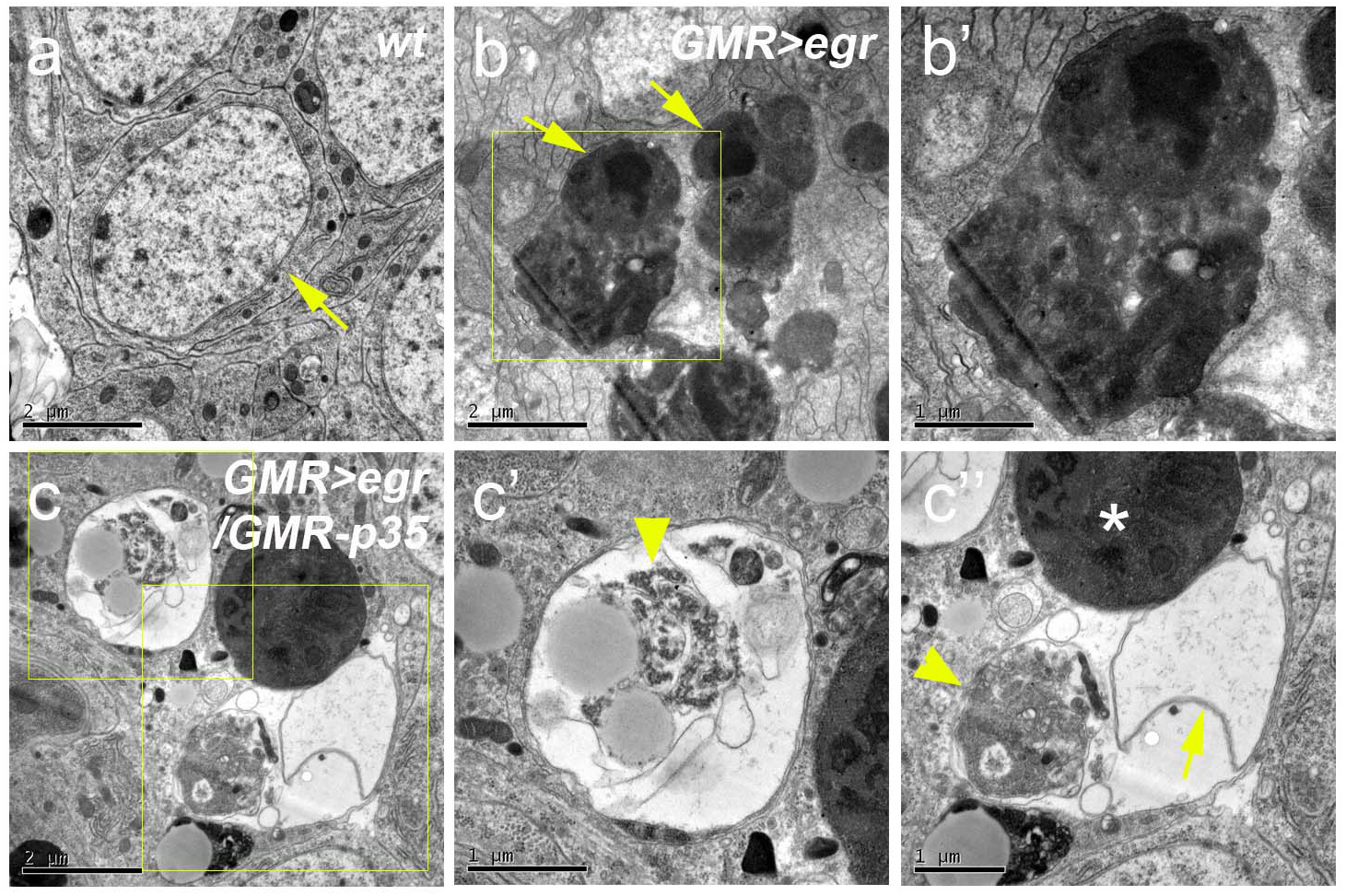
Non-apoptotic cell death in *GMR>egr/GMR-p35* shows morphological features of necrosis. TEM images of *wt* (a), *GMR>egr*/+ (b,b’) and *GMR>egr/GMR-p35* (c-c’’) discs at APF22h. b’ is an enlarged image for the outlined area in b. c’ and c’’ are enlarged images for the outlined areas in c. Compared to the wild type eye disc cell which has a large nucleus (yellow arrow, a), apoptotic features such as high-electron-density chromatin condensation (yellow arrows, b) and apoptotic bodies (dark aggregates in b and b’) are frequently observed in *GMR>egr* discs (b). Expression of P35 in *GMR>egr* (c), however, induces necrotic cell features such as translucent cytoplasm, mal-shaped (arrow, c’’) or unidentifiable nuclei (C’), and aggregation of endoplasmic reticulum and other cellular organelles (arrowheads, c’ and c’’). Asterisk indicates a phagolysosome.

### JNK signaling is required for Eiger-induced necrosis when apoptosis is blocked

Unlike *GMR>egr, hid*-induced apoptosis e.g. *GMR-hid* can be suppressed by expression of P35 (ref.^32^) or reduction of DrICE and Dcp-1 (Fig.4g-i). Therefore, factors other than inhibitors of apoptosis are also required for induction of necrosis in *GMR>egr/GMR-p35* discs. Because Egr activates JNK signaling upstream of *hid*-mediated apoptosis, we examined whether there is non-apoptotic input from the JNK signaling contributing to induction of necrosis. Hypomorphic mutants of *bsk, hep, MKK4* and *Tak1*,genes encoding various kinases in the JNK pathway ^53^, were used. We observed that heterozygosity of these mutants can only weakly or moderately suppress *GMR>egr*-induced eye ablation phenotype (compare Fig.7a-d to Fig.2b). Intriguingly, except *hep^1^* (Fig.7f), heterozygosity of other mutants including *bsk^1^, MKK4^G680^* and *Tak1^2^* can strongly suppress *GMR>egr/GMR-p35*-induced small eyes (compare Fig.7e,g,h to Fig.4d). This suggests that, in addition to activate apoptosis through *hid*, JNK signaling also contributes to induction of necrosis when apoptosis in blocked (Fig.7i).

**Figure 7.**
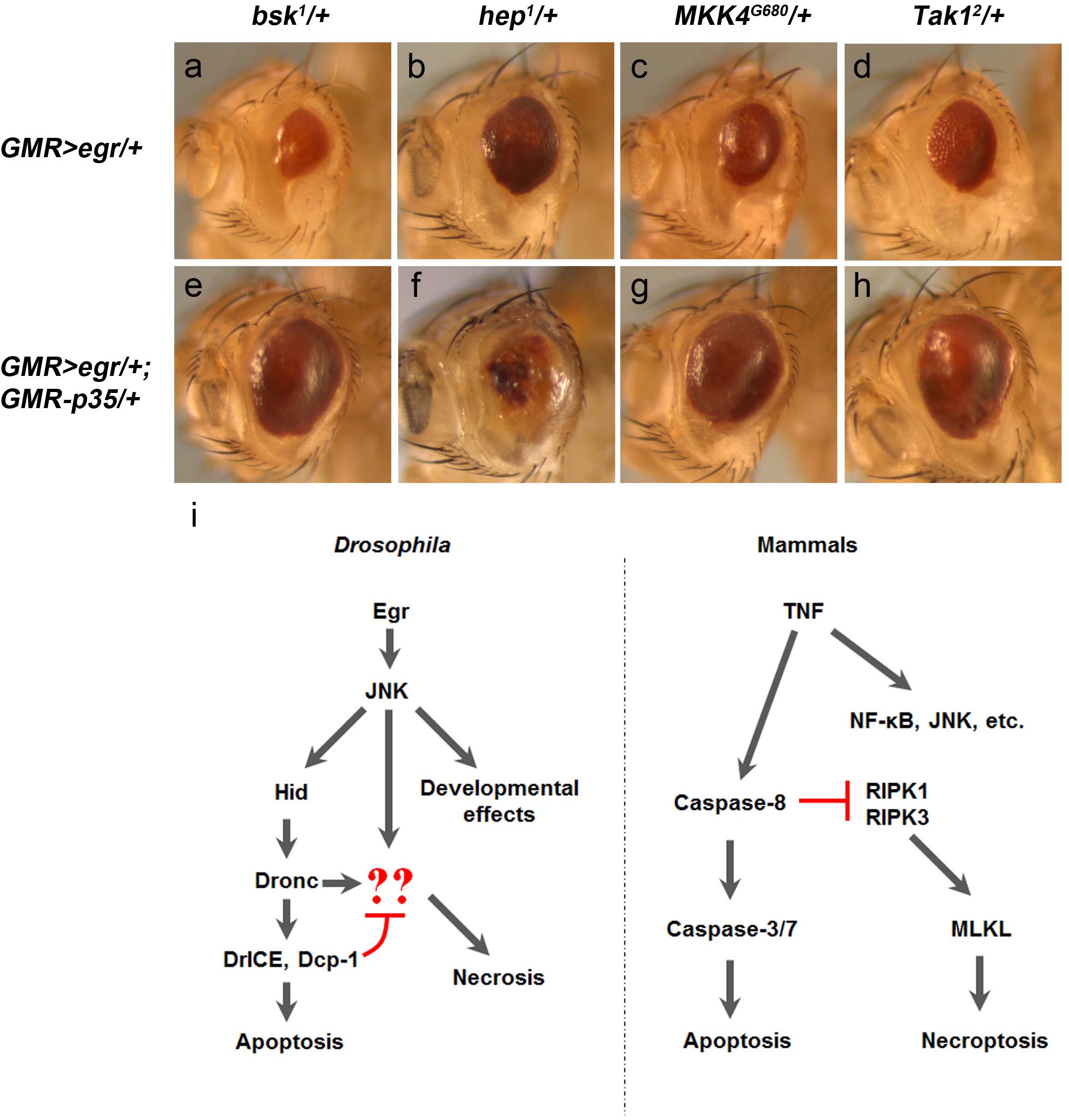
JNK signaling contributes to Eiger-induced necrosis when apoptosis is blocked. (a-h) Adult eye images. Heterozygous mutants of *bsk^1^*(a), *hep^1^*(b), *MKK4^G680^*(c) and *Takl^2^*(d) can only weakly or moderately suppress *GMR>egr*-induced eye ablation phenotype. In contrast, *GMR>egr/GMR-p35*-induced small eyes are strongly suppressed by heterozygous mutants of *bsk^1^*(e), *MKK4^G680^*(g) and *Takl^2^*(h), but not *hep^1^* (f). (i) A diagram showing comparable molecular mechanisms of regulated necrosis in *Drosophila* and mammals. *Drosophila* TNF (Egr), similar to its mammalian counterparts, has multiple context-dependent functions including induction of necrosis when apoptosis is blocked. In mammals, necroptosis (a type of regulated necrosis) can occur when inhibition of caspase-8 on RIPK1 and RIPK3 is removed. While in *Drosophila*, effector caspases DrICE and Dcp-1 inhibit necrosis. Once this inhibition is removed, the initiator caspase Dronc can activate necrosis with additional input(s) from JNK signaling. Key factors that mediate this necrosis downstream of caspases are currently unknown (question marks in red).

## Discussion

Our findings in this study suggest an analogy between *Drosophila* and mammals in regulation of TNF-induced necrosis (Fig.7i). Caspases play critical roles in these processes. In mammals, caspase-8 exerts a permissive role on apoptosis but an inhibitory role on necroptosis, a form of regulated necrosis. Here, we report that, in *Drosophila*, regulated necrosis can also occur when apoptosis is blocked. Interestingly, both apoptosis and necrosis induced by Egr, the *Drosophila* TNF, depend on the initiator caspase Dronc. However, loss of one copy of *dronc* blocks necrosis, but not apoptosis, suggesting that different levels of Dronc are required for apoptosis versus necrosis. In contrast to Dronc, effector caspases DrICE and Dcp-1, in particular DrICE, appear to inhibit Egr-induced necrosis. This inhibition can be released either by expression of the inhibitor P35 or reduction of DrICE and Dcp-1 by RNAi or mutants. However, inhibition of DrICE and Dcp-1 alone following apoptotic stresses is not sufficient to induce necrosis. For example, no necrosis is evident when apoptosis induced by expression of *hid* is inhibited (Fig.4g-i).^29^ This is further supported by our observation that heterozygous mutants of *bsk, MKK4* and *Tak1*, genes encoding various kinases in the JNK pathway, suppress eye ablation in *GMR>egr/GMR-p35* much better than in *GMR>egr*. Therefore, an additional input from Egr-induced JNK signaling is also required for induction of necrosis (Fig.7i).

How do DrICE and Dcp-1 inhibit Egr-induced necrosis? How does Dronc promote it when DrICE and Dcp-1 are inhibited? What factors coordinate the inputs from Dronc and JNK in this process? These are intriguing questions remained open (Fig.7i, question marks in red). In mammals, caspase-8 cleaves and inactivates RIPK1 and PIPK3 therefore inhibits TNF-induced necroptosis (Fig.7i). While pyroptosis, another type of regulated necrosis induced by pathogens, is activated by several inflammatory (non-apoptotic) caspases, such as caspase-1, –11 in mice and caspase-1, –4, –5 in humans, through their cleavage of gasdermin-D (GSDMD).^54^ Interestingly, capase-1, –4, –5 and –11 are structurally similar to the apoptotic initiator caspases. However, no homologs of RIPK1, RIPK3 and GSDMD have been reported in *Drosophila*. Our study suggests that the catalytic activity of Dronc is required for Egr-induced necrosis because only expression of the wild type Dronc, but not the catalytic site-mutated form of Dronc, can restore such necrosis in *dronc* mutants. Therefore, the key to understanding Egr-induced necrosis in *Drosophila* is to identify the non-apoptotic substrates of Dronc, DrICE and Dcp-1 that mediate the process. Interestingly, roles of necrosis-like cell death in *Drosophila* have also been reported in stress-induced loss of neural stem cells in the larval brain^52^, the development of male testes^55^ and female ovaries^56^. In these processes, the catalytic activity of Dronc or even Dronc itself doesn’t seem to be required. Hence, further studies are needed to understand how these different types of necrosis are regulated at the molecular level. Notably, a series of intracellular events specific to necrosis including mitochondrial dysfunction, ATP depletion, increased cytosolic Ca^2+^and organelle clustering appear to be conserved in multiple organisms in which necrosis has been studied.^57^ Are there conserved mechanisms regulating these necrotic cellular events throughout evolution despite death factors executing necrosis might be different across eukaryotes? Future genetic studies on various necrosis models in *Drosophila* will likely provide some insights into this question.

In addition to cell death, mammalian TNFs, as a family of pleiotropic cytokines, regulate a variety of cellular processes through activating multiple signaling pathways such as the NF-κB, JNK and p38-MAPK pathways (Fig.7i).^6, 10^ In *Drosophila*, although Egr exerts its functions mainly through the JNK pathway, context-dependent roles of the Egr-JNK signaling have also been reported in various cellular processes.^25^ Consistent with previous reports ^21, 22, 28^, we found that *GMR>egr* induces not only apoptotic but also non-apoptotic effects downstream of JNK activation in the developing *Drosophila* eye. The apoptotic effects are mediated by the pro-apoptotic gene *hid* and the canonical apoptosis pathway.

While the non-apoptotic effects are characterized by irregular ommatidial fusion and interommatidial spacing which may lead to the glassy appearance of the adult eyes. Recent studies suggest that these Egr-induced non-apoptotic effects might be mediated by the metabolic energy production pathways and the Toll/NF-κB pathway, a pathway regulates both immunity and development in *Drosophila*.^48, 58^ Further studies are required to elucidate how these pathways are coordinated to regulate eye development in response to Egr-induced activation of JNK signaling. It is also interesting to investigate whether these pathways contribute to Egr-induced necrosis when apoptosis is blocked.

## Material and Methods

### *Drosophila* genetics

Genetic crosses for all experiments were reared at 25°C. *GMR-GAL4 UAS-egr* (*GMR>egr*) ^22^, *GMR-hid^10^* (ref.^32^), *dcp-1^Prev1^* (ref.^59^), *drICE^Δ1^* (ref.^34^), *dronc^129^* (ref.^39^), *rpr^87^*(ref.^44^), *XR38* (ref.^43^), *hid^05014^* (ref.^32^), *GMR-BIR^41^, GMR-p35* (ref.^29^), *hid^20-10^-lacZ^42^, rpr^XRE^- lacZ^42^, sev>Glu^Lc50^, MKK4^G680^* (ref.^60^), *UAS-dronc^wt^* and *UAS-dronc^C>A 13^* were as described. *bsk^1^, hep^1^, Tak1^2527^, Tak1^2^* and *UAS-bsk^DN^* were obtained from the Bloomington stock center. *UAS-drICE^RNAi^* and *UAS-dcp-1^RNAi^* were obtained from the NIG-Fly stock center. *UAS-egr^RANi^* (108814) was obtained from the VDRC stock center. For mosaic analysis with *hid* mutant clones, larvae of the following genotype were analyzed at the late 3rd instar larval stage: *ey-FLP/+; *GMR>egr*/+; FRT80B /hid^05014^ FRT80B*.

### Immunohistochemistry

Pupal or larval discs were dissected, fixed (with 4% paraformaldehyde for 30min at room temperature), and then labeled with antibodies using standard protocols. Primary antibodies used are rabbit anti-cDcp1 (the cleaved Dcp-1 antibody, 1:500, Cell Signaling), rat anti-ELAV, mouse anti-Dlg and mouse anti-β-Gal (all 1:50, DHSB). Secondary antibodies were goat Fab fragments conjugated to Alex488, 555, or 647 (all 1:1000) from Molecular Probes.

### Propidium Iodidum (PI), Hoechst labeling and TUNEL

For PI and Hoechst double labelling on pupal discs, freshly dissected discs were incubated in dark with 4μM PI (Sigma-Aldrich) and 16μM Hoechst (ThermoFisher) in Schneider’s media for 1h at room temperature as previously described.^51^ The discs were then fixed with 4% paraformaldehyde for 20min at room temperature followed by gentle washes. For PI labeling alone, larval or pupal eye discs were incubated with 15 μI for 10min at room temperature followed by fixation (with 4% paraformaldehyde for 20min at room temperature) and gentle washes as previously reported.^50^ PI labeling can also be observed and scored without fixation. For TUNEL (terminal deoxynucleotide transferase-mediated dUTP end labelling), dissected and fixed (with 4% paraformaldehyde for 20min at room temperature) discs were incubated in 100 mM Na-citrate with 0.1% Triton X-100 for 30 min at 65°C, followed by detection of dying cells using an in situ cell death detection kit (Roche).

### Imaging and statistical analysis

Fluorescent eye disc images were taken with either a Zeiss or Leica confocal microscope. Adult fly eye images were taken using a Zeiss stereomicroscope equipped with an AxioCam ICC1 camera. For statistical analysis of PI-labeling (Fig.5c), at least 20 pupal discs from each genotype were used for counting the number of PI-positive cells. For quantification of adult eye size (Fig.5d), the ‘‘histogram’’ function in Adobe Photoshop CS was used to measure at least 10 representative adult eyes of each genotype. The statistical significance was evaluated through a one-way ANOVA with Bonferroni multiple comparison test using GraphPad Prism.

### Transmission electron microscopy (TEM)

Freshly dissected pupal eye-brain complexes were fixed in 2.5% glutaraldehyde in 0.1M sodium cacodylate buffer (pH 7.4) for 45 min followed by a secondary fixation of 1% Osmium Tetroxide for 1 hour at room temperature. The samples were then washed with the buffer (10×5min) and dehydrated in ascending concentrations of ethanol before they were embedded in epoxy resin. Sections (90nm) were prepared and stained with uranyl acetate and lead citrate followed by examination using a JEOL 1200EX electron microscope fitted with a tungsten filament. Images were acquired through a GATAN MultiScan camera.

## Acknowledgement

We apologize to colleagues whose work was not cited due to space limitations. We would like to thank Paul Stanley and Theresa Morris at the Centre for Electron Microscopy in the University of Birmingham for their assistance on TEM. We thank Kai Liu for advice on PI-labeling. We are grateful to Konrad Basler, Andreas Bergmann, Lei Liu, Pascal Meier, the Bloomington Stock Center, the NIG-Fly stock center in Japan, the VDRC stock center in Vienna and the Developmental Studies Hybridoma Bank (DSHB) in Iowa for fly stocks and reagents. ML is supported by the China Scholarship Council (CSC)-Birmingham joint PhD program. YF is supported by the Birmingham Fellowship, Grant BB/M010880/1 from the Biotechnology and Biological Sciences Research Council (BBSRC) UK and the Marie Curie Career Integration Grant (CIG) 630846 from the European Union’s Seventh Framework Programme (FP7).

## Conflict of Interest

The authors declare that there is no conflict of interest.

## Supplemental Figures S1-S3

**Supplemental Figure S1.*GMR>egr* induces transcription of the pro-apoptotic gene *hid*.**(a-b’) Expression of *hid-lacZ* (indicated by βGAL, red in a, b and grey in a’, b’), a reporter of transcription of *hid*, and cDcp1 labeling (green in a,b) in late 3^rd^ instar larval discs, anterior is to the left. Compared to the control (a,a’), expression of *hid-lacZ* is moderately induced by *GMR>egr* in the *GMR* domain (yellow outlined, a’ and b’) where Egr is expressed (b,b’).

(c) Quantification of *hid* expression levels as indicated by normalizing βGAL signal intensity in the *GMR* domain (yellow outlined, a’ and b’) to the area in the antenna (blue outlined, a’ and b’) where *GMR* is not expressed. Increase of *hid* expression in *GMR>egr* is statistically significant (P<0.001, Student’s t-test).

(d-e’) Expression of *rpr-lacZ* (indicated by βGAL, red in d, e and grey in d’, e’), a reporter of transcription of *rpr*, and cDcp1 labeling (green in d,e) in late 3^rd^ instar larval discs, anterior is to the left. Compared to the control (d,d’), expression of *rpr-lacZ* does not increase in *GMR>egr* in the *GMR* domain (yellow outlined, d’ and e’) where Egr is expressed (e,e’).

(f) Quantification of *rpr* expression levels as indicated by normalizing βGAL signal intensity in the *GMR* domain (yellow outlined, d’ and e’) to the area in the antenna (blue outlined, d’ and e’) where *GMR* is not expressed. No significant increase (n.s.) of βGAL signal intensity in *GMR>egr* was detected.

**Supplemental Figure S2.cDcp-1 labels developmental apoptosis**. Pupal discs at APF28h labeled with cDcp1 (green), TUNEL (red) and ELAV (blue). Strong developmental apoptosis as indicated by cDcp1 and TUNEL occur at APF28h in wild type discs (a-a’’). This developmental apoptosis is completely suppressed by expression of P35 (b-b’’).

**Supplemental Figure S3.No PI-labeling was detected in *GMR>egr/GMR-p35* larval eye discs**. Late 3^rd^ instar larval eye discs labeled with Propidium Iodide (PI). In contrast to *Sev>Glu^*LC*^*(b) in which necrosis, indicated by Pi-positive cells, is induced (ref. S1), no PI labeling were detected in wild type (a), *GMR>egr* (c) or *GMR>egr/GMR-p35* (d) larval eye discs.

